# Defining kinetic roles of transcriptional activators in the early Drosophila embryo

**DOI:** 10.1101/2021.02.25.432925

**Authors:** Timothy T. Harden, Ben J. Vincent, Angela H. DePace

## Abstract

Most animal transcription factors are categorized as activators or repressors without specifying their mechanisms of action. Defining their specific roles is critical for deciphering the logic of transcriptional regulation and predicting the function of regulatory sequences. Here, we define the kinetic roles of three activating transcription factors in the Drosophila embryo—Zelda, Bicoid and Stat92E—by introducing their binding sites into the *even skipped* stripe 2 enhancer and measuring transcriptional output with live imaging. We find that these transcription factors act on different subsets of kinetic parameters, and these subsets can change over the course of nuclear cycle (NC) 14. These transcription factors all increase the fraction of active nuclei. Zelda dramatically shortens the time interval between the start of NC 14 and initial activation, and Stat92E increases the duration of active transcription intervals throughout NC 14. Zelda also decreases the time intervals between instances of active transcription early in NC 14, while Stat92E does so later. Different transcription factors therefore play distinct kinetic roles in activating transcription; this has consequences for understanding both regulatory DNA sequences as well as the biochemical function of transcription factors.

## INTRODUCTION

In all cells, gene transcription is activated or repressed by transcription factor proteins (TFs) which bind DNA in a sequence-specific manner and are accessory to the molecular machinery that is required to carry out transcription. Some TFs downregulate transcription—repressors—while other TFs upregulate transcription—activators (Johnson, 1995; Workman and Buchman, 1993). These labels assign individual proteins to broad classes of functional activities. Since the advent of this activator/repressor paradigm (Hughes, 2011), functional subclasses of activators and repressors have been delineated by assigning more specific mechanistic labels to individual TFs. For example, pioneer factors open local chromatin allowing subsequent binding of other factors (Zaret and Carroll, 2011); short- and long-range repressors work to silence nearby or distally bound activators, respectively (Courey and Jia, 2001; Gray and Levine, 1996); and bifunctional factors exhibit context-dependent activity with the capacity to either activate or repress transcription (e.g. (Majello et al., 1997; Staller et al., 2015; Stampfel et al., 2015)).

Aside from a handful of notable exceptions (e.g. (Duarte et al., 2016)), most animal TFs remain categorized as activators, repressors, or bifunctional factors (Lambert et al., 2018). This stands in contrast to bacteria, where the activator/repressor paradigm is rich with detailed descriptions of TF mechanisms (Browning and Busby, 2016). There, biochemical and structural approaches have elucidated detailed chemical and physical mechanisms for many individual TFs (e.g. the sigma factors (Bae et al., 2015; Chen et al., 2020; Friedman and Gelles, 2012; Harden et al., 2016)). Within animal transcription, research has largely focused on a tissue-specific paradigm of TF function that identifies TFs responsible for developmental patterning and cell type specification and characterizes them as activators or repressors (Guo et al., 2018; Lambert et al., 2018; Villar et al., 2014). The stage is thus set for the animal activator/repressor paradigm to be fleshed out in greater detail (Lis, 2019).

Mechanistic information on TF function has typically been obtained using biochemistry and fluorescence imaging. The in vitro reconstitution approaches that have proven indispensable in the study of bacterial transcription regulation are transferable to eukaryotic model organisms (Fazal et al., 2015; Rosen et al., 2020), yet remain challenging (Chen and Larson, 2016). In vivo detection of nascent transcript synthesis via the MS2/MS2 coat protein (MCP) system has emerged as the technique of choice for studying transcription regulation at the level of individual genes in eukaryotes, and specifically in *Drosophila melanogaster* embryos (Bentovim et al., 2017; Lenstra et al., 2016; Lim, 2018). This system has been used to measure activation by individual TF proteins in the fly embryo by either quantifying changes in TF concentration (Bothma et al., 2018; Eck et al., 2020) or through mutation of regulatory DNA to introduce or disrupt TF binding sites (Dufourt et al., 2018; Keller et al., 2020; Yamada et al., 2019).

For MS2/MCP experiments, the most challenging part of the technique is no longer making the measurements, but rather analyzing the resulting data and deriving mechanistic conclusions from it. Many studies have measured transcription in the embryo using the MS2/MCP system (reviewed in (Fernandez and Lagha, 2019; Wissink et al., 2019)).The analytical approaches employed by these studies range from statistical quantification (e.g. (Fukaya et al., 2016; Yamada et al., 2019)) to various mathematical models (Bothma et al., 2015; Desponds et al., 2016, 2020; Dufourt et al., 2018; Eck et al., 2020; Keller et al., 2020; Lammers et al., 2020). However, the MS2/MCP measurements themselves are many biochemical steps removed from the molecular kinetics of interest, namely transcription initiation. This makes the application of predictive models derived from biochemical pathways (e.g. (Lammers et al., 2020)) difficult. Here, we analyze MS2/MCP data with approaches used to elucidate kinetics from single molecule in vitro transcription experiments (Friedman and Gelles, 2012). This empirical approach does not predict transcriptional outputs a priori and cannot elucidate the kinetics of the biochemical steps that lead to transcription. Instead, it gives insight into the function of TFs by tracking changes in kinetic parameters—the timing and duration of MS2/MCP signal—due to increased activator activity.

We characterized the kinetic mechanisms of three activator proteins present in the early *D. melanogaster* embryo. Zelda (Zld) is uniformly distributed across the early embryo (Staudt et al., 2006) and is thought to be a pioneer factor that can establish and/or maintain open chromatin ((Harrison and Eisen, 2015) and references therein). Zld has been previously shown to decrease the time of first passage into transcription within the blastoderm (Dufourt et al., 2018; Eck et al., 2020). Bicoid (Bcd) is a *Hox3*-derived protein that is well known for its role in patterning the anterior-posterior axis of the embryo through a concentration gradient (McGregor, 2005; Struhl et al., 1989) and is dependent upon inter-protein cooperative interactions to activate transcription (Burz et al., 1998; Ma et al., 1996, 1999; Park et al., 2019). Stat92E (Dstat) is the signal transducer and activator of transcription (STAT) component in the *Drosophila* JAK/STAT pathway (Herrera and Bach, 2019). Dstat is uniformly distributed across the early embryo, is an essential zygotic activator (Tsurumi et al., 2011; Yan et al., 1996), and has been proposed to act downstream of nucleosome displacement to activate transcription (Barr et al., 2017).

To decipher the kinetic function of Zld, Bcd, and Dstat, we created transcription reporters containing the *even-skipped* stripe 2 minimal enhancer (*eve2*) and its cognate promoter (Small et al., 1992). Activation through *eve2* has been highly studied, both in terms of the cis-regulatory sequences required (Arnosti et al., 1996; Goto et al., 1989; Small et al., 1992; Vincent et al., 2016) and the spatiotemporal pattern it drives (Berrocal et al., 2020; Bothma et al., 2014; Lim et al., 2018). This makes *eve2* an ideal substrate for this detailed kinetic study.

In this study, we used MS2 transcription reporters and kinetic models to determine the roles of three individual transcriptional activators. Transcriptional dynamics driven by variants of *eve2* containing additional binding motif sequences for Zld, Bcd, and Dsat were compared to the dynamics of two benchmark sequences: a 1.5 kb 5′ flanking region of the endogenous *even-skipped* locus including *eve2*, and a minimal regulatory sequence, where *eve2* is separated from its promoter by 765 bp of synthetic spacer sequence which is not predicted to bind any TFs present in the early embryo. We then applied a collection of chemical kinetics-based models to characterize transcription and its regulation driven by these sequence variants. These models provided a set of kinetic parameters that were used to determine the mechanism of regulation for the TFs. We found that Zld, Bcd, and Dstat each acted on overlapping but unique subsets of parameters over the course of nuclear division cycle (NC) 14. The kinetic role of each TF also changed over the course of the nuclear cycle. This work provides a framework for functionally classifying transcriptional activators by the kinetic parameters they modulate and yields insight into the combinatorial control of transcription.

## RESULTS

### *eve2* separated from the promoter drives weak expression

To investigate the kinetic roles of individual activating TFs, we sought a transcription reporter that would give mechanistic insights into the individual proteins that bind a regulatory sequence. Activation by the *eve2* enhancer is driven by binding sequence motifs for Bcd (Small et al., 1991), along with additional sequence elements that are not exactly known (Barr et al., 2017; Vincent et al., 2016, 2018), but likely include those for Dstat (Barr and Reinitz, 2017) and Zld (Crocker and Stern, 2017; Struffi et al., 2011). The anterior and posterior edges of the stripe 2 domain are set by repressor proteins, including Giant and Kruppel, that bind to sequences within *eve2* (Small et al., 1992). *Dynamic expression driven by eve2* has been previously measured in two contexts: the endogenous *even-skipped* locus (Berrocal et al., 2020; Lim et al., 2018) and a reporter for 1.7 kb of the *even-skipped* 5′ flanking region (Bothma et al., 2014). To isolate regulation from *eve2* alone (rather than its flanking sequences), we constructed a reporter, *eve2:neutral*, containing *eve2* and the *even-skipped* promoter separated by a 765 bp neutral sequence spacer (Fig. 1A). The spacer sequence was computationally designed to lack predicted binding sites for transcription factors present in the early embryo (Estrada et al., 2016) (see Methods). The spacer length is comparable, though not identical, to the distance between eve2 and the eve promoter at the endogenous locus (1033 bp). We did not place the enhancer immediately upstream of the promoter because TFs bound to enhancers placed immediately adjacent to the promoter can act differently than they do when placed at a distance (Scholes et al., 2019; Weingarten-Gabbay and Segal, 2014). The reporter contained 24 tandem repeats of an MS2 binding motif sequence (MBS) (Hocine et al., 2013) in the 5′ untranslated region of a transcription unit (see Methods) and was integrated into the attP2 landing pad site using phiC31-mediated transgenesis (Groth et al., 2004).

**Figure 1.**
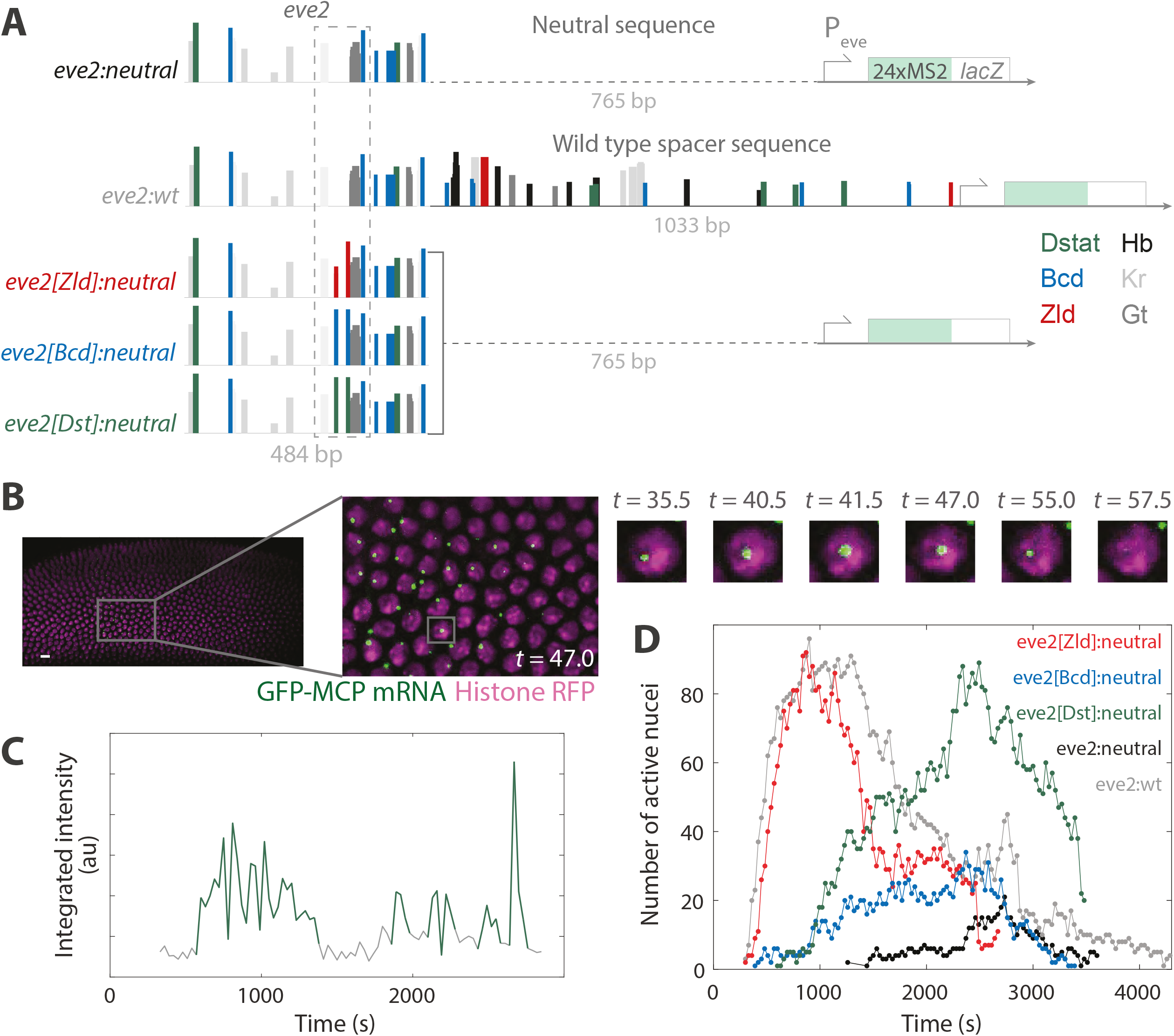
Measuring the activity of individual TFs against benchmark regulatory sequences. (A) Schematics of the minimal *even-skipped* stripe two enhancer (*eve2*) transcription reporter constructs. Each contains the *even-skipped* promoter driving expression of 24 repeats of the MS2 stem loop sequence followed by a partial sequence of the bacterial *lacZ* operon. *eve2:neutral* and *eve2:wt* are benchmark reporters with which we compare all others to. *eve2:neutral* contains a spacer sequence with no predicted transcription factor binding sites (dashed line). *eve2:wt* is *eve2* with a spacer containing the wild type locus sequence between the enhancer and the promoter. The three transcription activator-specific reporters*—eve2[Zld]:neutral, eve2[Bcd]:neutral, and eve2[Dst]:neutral—*are identical to *eve2:neutral* but contain two mutations to add predicted binding motifs for a single transcription activator (dashed box), either Zld, Bcd, or Dstat, respectively. Colored bars are transcription factor binding sites predicted by the software SiteOut (Estrada et al., 2016). Hb, Kr, and Gt stand for Hunchback, Kruppel, and Giant, respectively. (B) Left: image of the diSPIM microscope field of view with histone-red fluorescent protein (magenta) and GFP-MS2 coat protein (green). Gallery images: magnified view of the marked region over time showing a detected active transcription locus. *t* = 0 corresponds to the beginning of nuclear cycle 14; time is in minutes. Scale bar 10 μm. (C) Example MCP-GFP fluorescence emission record from a single nucleus during NC 14. Green marks detected active transcription signal; gray marks intervals during which no fluorescent signal was detected. (D) Dynamic transcription profiles during NC 14 for the constructs in (A). Binary detection of the number of detected active transcription loci within the microscope field of view from two replicate experiments for each condition. *t* = 0 is the end of anaphase.

Living embryos were imaged by dual-inverted selective plane illumination microscopy (diSPIM) (Wu et al., 2013). Previous studies have used confocal microscopy (Dufourt et al., 2018; Fukaya et al., 2016; Garcia et al., 2013; Lucas et al., 2013; Scholes et al., 2019; Zoller et al., 2018), which relies on oil-immersion objective lenses and requires subjecting the embryos to continuous submersion in halocarbon oil before and during imaging. In contrast, diSPIM relies on water-immersion objective lenses, allowing dechorionated embryos to be placed on a single coverslip and imaged while in a bath of Schneider’s media (Methods). We found that embryos were equally viable when imaged using both techniques (Scholes et al., 2019). However diSPIM has other advantages, namely decreased photobleaching rates and phototoxicity for a comparable signal-to-noise ratio (Icha et al., 2017; Jemielita et al., 2013; Laissue et al., 2017), without loss of spatial or temporal resolution.

We observed the appearance of MS2 coat protein-green fluorescent protein (MCP-GFP) signal as diffraction-limited spots above background (Fig. 1B). This signal was colocalized with histone-red fluorescent protein (his-RFP) signal, reflecting the binding of many MCP-GFP proteins to nascent RNA in complex with actively transcribing RNA polymerase II proteins (pol II) within individual nuclei (Bertrand et al., 1998). As in previous studies, we interpreted the appearance of signal from the MBS repeats as the start of active transcription (Fukaya et al., 2017; Garcia et al., 2013; Lucas et al., 2013). When the transcription signal later disappeared, presumably due to transcript termination by most or all actively transcribing pol II and subsequent release of the fluorescently tagged mRNA, this was scored as the end of active transcription. Oftentimes, we observed multiple instances of transcription activation within the same nucleus over the time course of NC 14 (e.g. Fig. 1C, green), consistent with a previous study of *eve2* (Bothma et al., 2014). Active transcription was not observed outside the stripe 2 pattern during NC 14, except in select cases, wherein transcript signal was detected within the domain of eve stripe 7, which was expected given previous reports that used transcription reporters for eve2 (Bothma et al., 2014; Janssens et al., 2006; Staller et al., 2015; Vincent et al., 2018). We limited our analysis to the nuclei located in the center of the stripe (SFig. 1), where repressive interactions by Giant and Kruppel are minimized (Barr and Reinitz, 2017; Small et al., 1992), in an attempt to isolate activating TF activity from repressive TF activity. For each imaging replicate, acquisition began during nuclear division cycle (NC) 13 and analysis was performed on all time points from the start of NC 14, defined here as the end of anaphase, until active transcription ceased with the onset of gastrulation. In a comparison of replicates, we found they were indistinguishable (SFig. 2). Analysis was conducted with custom Matlab software that was adapted from an existing platform and based on signal detection and tracking methods previously used to analyze single molecule fluorescence data ((Friedman and Gelles, 2015); see Methods).

We observed weak expression driven by eve2:neutral. No nuclei exhibited MCP-GFP signal above detection threshold until ∼20 min into NC 14 (Fig. 1D, black curve), compared to <10 min reported by Bothma et al. for an MS2 reporter driven by eve2 flanked by sequences from the endogenous even-skipped locus. In addition, transcription driven by eve2:neutral was detected in a small number of nuclei compared to that same reporter (Bothma et al., 2014).

### An extended region upstream of *even-skipped* which includes *eve2* drives a normal pattern of expression

To investigate the cause of the weak expression driven by *eve2:neutral*, we created a second reporter, *eve2:wt*, containing a 1517 bp sequence identical to the 5′ flanking region of the endogenous *even-skipped* locus. This sequence was composed of *eve2* and a 1033 bp spacer sequence—which contains additional regulatory elements—between *eve2* and the promoter (Fig. 1A). An MS2 reporter with a similar extended version of *eve2* was previously shown to drive a normal pattern of expression (Bothma et al., 2014). *eve2:wt* was also integrated into attP2, and imaged as described above.

We observed a stark difference in transcription driven by *eve2:wt* compared to that of *eve2:neutral* throughout NC 14 (Fig. 1D, gray curve). Transcription driven by *eve2:wt* was detected earlier in the nuclear cycle, occurred in more nuclei, and was more persistent throughout NC 14.

### Designing regulatory sequences to measure the activity of individual TFs

Ultimately, we sought to measure the roles of different activating TFs. This required a regulatory context in which the consequences of additional activation are not obscured by a high baseline of transcription signal; in previous studies, high levels of activity were thought to obscure the detection of transcriptional bursts (Fukaya et al., 2016; Garcia et al., 2013; Lucas et al., 2013). The *eve2:neutral* reporter, with its weak basal expression, provides this context. We therefore created a set of variants of *eve2:neutral* designed to recruit additional specific TFs to the transcription locus. Within the bounds of *eve2*, each of these variants contained two short (8 to 10 nucleotides) sequence mutations to introduce two DNA binding motifs for a single TF—either Zld, Bcd, or Dstat (See Methods). Each of these activating TF reporters—*eve2[Zld]:neutral, eve2[Bcd]:neutral*, and *eve2[Dst]:neutral*—contained the same neutral sequence spacer as *eve2:neutral*, and were incorporated into attP2 and imaged as above (Fig. 1A). In contrast to *eve2:neutral, eve2:wt*, which contained additional sequences that encode predicted binding motifs for Zld, Bcd, and Dstat, drove early, persistent transcription (Fig. 1D) in nearly all the nuclei within the center of the stripe 2 domain (SFig 1). Therefore, we treated *eve2:wt* as an effective upper-bound benchmark condition to compare with *eve2[Zld]:neutral, eve2[Bcd]:neutral*, and *eve2[Dst]:neutral*.

All three of these activating TF reporters induced transcription that exceeded that of *eve2:neutral*. Each dramatically altered the dynamic transcription profile (Fig. 1D) and did so in a unique way. *eve2[Bcd]:neutral* induced transcription similar to that of *eve2:neutral*, but earlier and in a greater number of nuclei. *eve2[Dst]:neutral* and *eve2[Zld]:neutral* also induced transcription earlier than *eve2:neutral* but in far more nuclei, similar to that of *eve2:wt*. However, the timing of activation by *eve2[Dst]:neutral* and *eve2[Zld]:neutral* was different. The dynamic transcription profile of *eve2[Zld]:neutral* peaked early in the nuclear cycle then decayed quickly, again similar to *eve2:wt*, while that of *eve2[Dst]:neutral* had a later, broader peak.

### Kinetic models distinguish the roles of Zld, Bcd and Dstat in regulating transcription

Elucidating kinetic mechanisms from dynamical data requires a mathematical model (Zhou and Zhuang, 2007). Mathematical models provide a modality to objectively compare conditions, rigorously interrogate the data, and have the potential to realize the rate limiting steps of regulation and to assign biochemical mechanisms to those steps (e.g. (Duarte et al., 2016)). The set of activating TF reporters, in comparison with the benchmarks *eve2:neutral* and *eve2:wt*, permits us to dissect the kinetic mechanisms that lead to increased transcription by each of these activating TFs.

The number and affinity of binding sites for each of Zld, Bcd, and Dstat differs within *eve2*, making it impossible to directly compare between activating TF reporters; instead, comparisons must be made to the benchmark conditions, either *eve2:neutral* or *eve2:wt*. To make these comparisons, we report the distributions of three different measurements—the first passage time, the active transcription lifetime, and the idle period—which we explain below. These MS2/MCP measurements are straightforward and their distributions are simple to extract from any live imaging dataset (see Discussion). From these, we determined kinetic parameters using models commonly used in the analysis of single molecule data (Friedman and Gelles, 2015).

#### First passage times

There are dramatic differences in the early time points of the dynamic transcription profiles of Fig. 1D. These differences are best represented in the distributions of times when transcription is first detected in each nucleus in NC 14—the first passage time (Fig. 2). These distributions have three important features that capture the mechanisms that lead to first passage into transcription. First, the maximum slope of the distribution is related to the rate of initial transcription activation across the stripe. Second, the plateau of the distribution is related to the fraction of active nuclei within the stripe. Third, the time delay between the end of anaphase (i.e. *t* = 0 in Fig. 2B) and the first detection of transcription within the stripe 2 domain (e.g. Fig. 2B *t* = 300 s and *t* = 1260 s for *eve2:wt* and *eve2:neutral*, respectively) is related to the number and length of kinetic states that a locus must pass through prior to reaching a state capable of initiating transcription.

**Figure 2.**
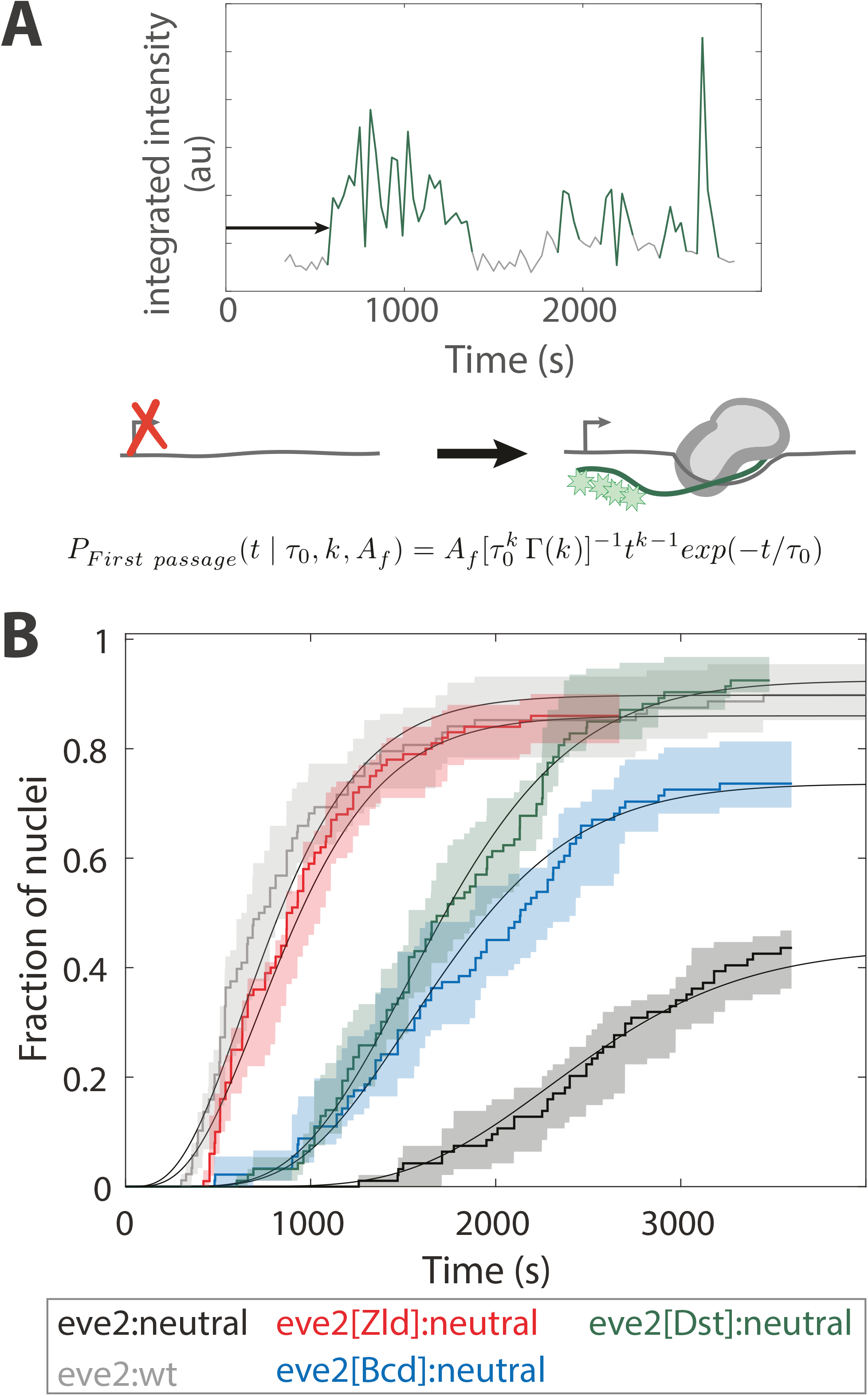
First passage activation kinetics. (A) Graphical depiction of a first passage transcription time measurement. Emission record as in Fig. 1C. The arrow denotes the first passage time for this nucleus. Cartoon: during this interval the transcription reporter transitions from a state incapable of activating transcription (left) to a transcriptionally competent one that activates transcription (right). The mathematical model (Eq. 1) is shown at the bottom. (B) Cumulative first passage distributions (stair-like curves) overlaid with a model (Eq. 1, smooth curves) with the characteristic number of rate limiting steps,*k*, a characteristic time constant for each of those steps, *τ*_0_, and the fraction of active nuclei within the center of the stripe,*A*_*f*_; see text for details on the interpretation of the model and Table 1 for parameter values. The curves are normalized to the total number of nuclei in the center of the stripe (see SFig. 1). Shaded regions represent the 90% confidence intervals from bootstrapping methods (see Methods).

Choosing a model to describe the first passage distributions has been challenging (Desponds et al., 2020; Dufourt et al., 2018; Eck et al., 2020). One approach has been to ignore the time delay following the end of anaphase and only consider the first passage times once transcription has been detected in any nucleus across the entire pattern (as in (Dufourt et al., 2018)). This is reasonable since transcription cannot take place during mitosis—a process called mitotic repression (Esposito et al., 2016; Parsons and Spencer, 1997). We initially ignored the time delay and attempted to apply the same models that are used to describe single molecule kinetics (Friedman and Gelles, 2015). These models performed poorly, as the observed time delay following anaphase requires a reaction path with more than two transcriptionally silent, rate limiting steps (explained in SFig. 3). Furthermore, the time delay varies by ∼900 s between the five different reporters (Fig. 2B), which suggests that TFs are acting to shorten the time delay (see Discussion). This means that the time delay, in general, cannot entirely be attributed to mitotic repression and thus cannot be ignored. We therefore employed a model that includes multiple transcriptionally silent kinetic steps; the possible biochemical mechanisms underlying these steps are described in the Discussion.

**Figure 3.**
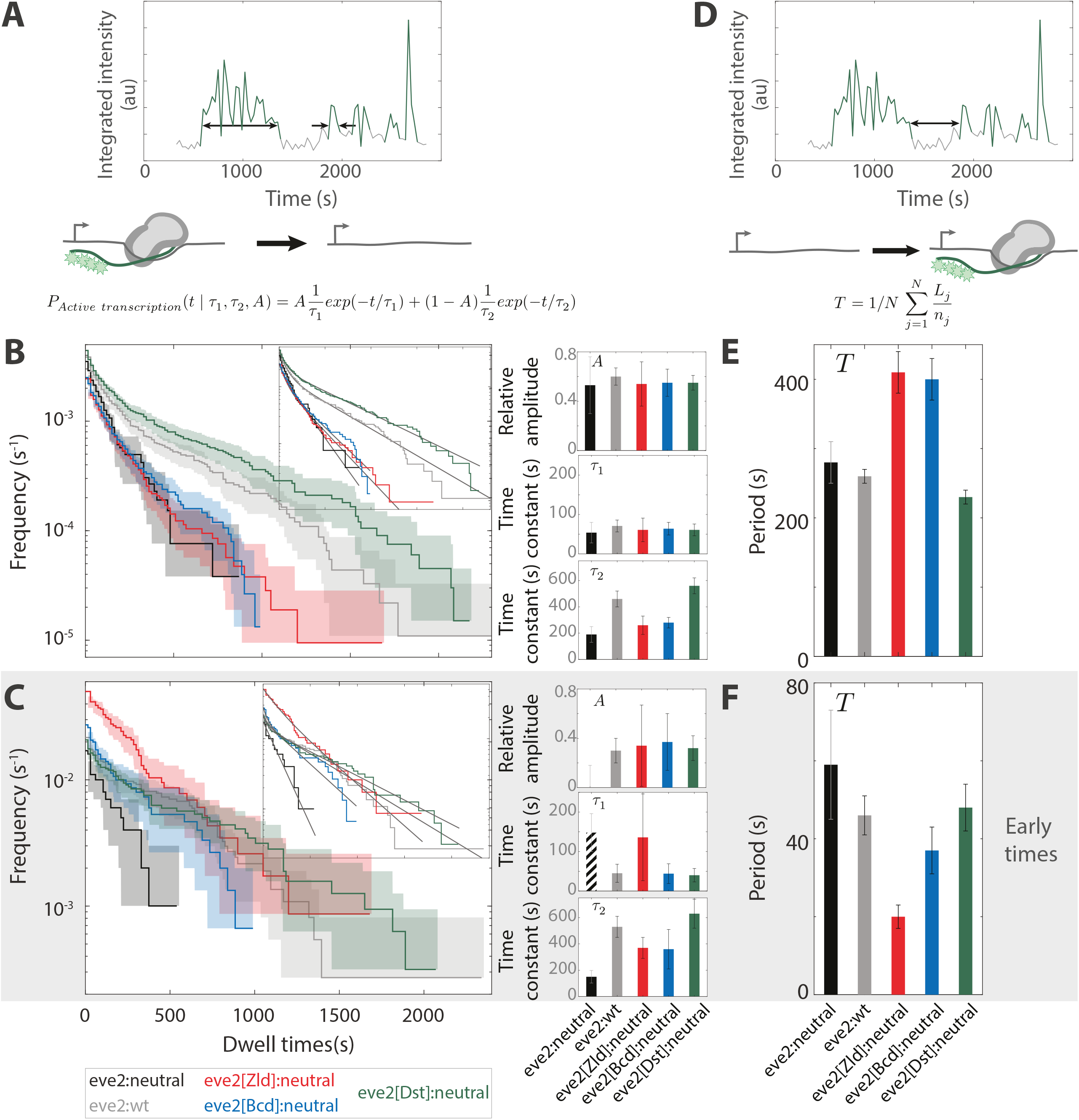
Active transcription and idle period kinetics. (A) Graphical depiction of active transcription lifetime measurements. Emission record as in Fig. 1C. The double headed arrows denote two example intervals of active transcription. Cartoon: during these intervals the transcription reporter locus transitions from a state containing many RNA polymerase molecules (gray bean) undergoing RNA synthesis (green line) to one lacking detectable active transcription (green stars). The model (Eq. 2) is shown at the bottom. (B) Cumulative frequency distributions of active transcription lifetimes (stair-like curves). The vertical axis is the average frequency with which active transcription intervals longer than the given interval lifetime (horizontal axis) occur. Plotting the data in this way gives the mean idle period as the vertical axis intercept. Data is overlaid with a model (Eq. 2; smooth curves) with time constants 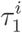 and 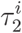 and relative amplitude *A*^*i*^, shown in bar charts to the right. Shaded regions represent the 90% confidence intervals from bootstrapping methods. (C) As in (B), but the data have been partitioned into the 20% of active transcription lifetimes that first appear during NC 14 (see Methods). (D—F). Idle transcription period. (D) Graphical depiction of idle transcription period measurements. Emission record as in Fig. 1C. The double headed arrow denotes a representative idle period. Cartoon: during these intervals the transcription reporter locus transitions from a state lacking active transcription to one containing many RNA polymerase molecules undergoing RNA synthesis. The model (Eq. 3) is shown at the bottom. (E) Barchart of mean idle periods, *T*, computed using Eq. 3; see Table 3 for parameter values. (F) As in (E), but for the 20% of idle periods that first occur during NC 14. All barcharts show standard errors from bootstrapping methods.

A critical choice for implementing this type of model concerns the characteristic time constants of each transcriptionally silent step. Although it is unlikely that they are all equivalent, it is reasonable that they are each of the same order of magnitude (Rosen et al., 2020; Scholes et al., 2016). We chose to assume the time constant of each silent step is equal, in alignment with previous studies (Dufourt et al., 2018; Eck et al., 2020). From this, we are forced to assume that the number of rate limiting steps must change to account for the differences in the first passage distributions in Fig. 2B. This is not to say that the kinetic pathway itself changes, only that under different circumstances some steps become fast and are no longer rate limiting. Finally, due to the limitations of the perturbations here, the model must be agnostic to exactly what the rate limiting steps represent biochemically and the order in which they occur.

From these choices, we arrived at a linear kinetic model of several steps, each with an equivalent characteristic time constant. Each step is assumed to be irreversible. This simplifying assumption is likely not appropriate for every step in the kinetic pathway, but, mathematically, any linear kinetic scheme containing reversible steps can be substituted by an equivalent scheme of irreversible steps through the addition of pseudo-steps that do not represent biochemical reactions (Scholes et al., 2016); the qualitative conclusions here will likely be unchanged even if the true kinetic scheme contains reversible steps. We chose to describe the first passage distributions with a gamma distribution model (a generalization of the poisson distribution):

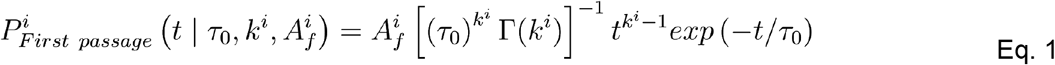

Where *i* ≡ *eve2:wt, eve2:neutral, eve2[Zld]:neutral, eve2[Bcd]:neutral*, or *eve2[Dst]:neutral*. 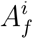 is the active fraction of nuclei in the center of the stripe, τ_0_ and *k*^*i*^ are the gamma distribution scale parameter and shape parameter, respectively, and Γ is the gamma function (Fig. 2B, smooth curves; Table 1). As was laid out in Dufourt et al, *k*^*i*^ can be interpreted as the average number of rate-limiting steps while τ_0_ can be interpreted as the characteristic time constant, τ_0_, of each of these steps. The time constant,, was fit globally to all five reporter data sets while, simultaneously, the number of steps, *k*^*i*^, was fit independently for each reporter.

**Table 1.** First passage into active transcription model parameters. See Eq. 1. The characteristic lifetime for all rate limiting steps, *τ*_0_, was globally fit to all five transgenic reporters. Standard errors were computed using bootstrapping methods (see Methods).

| | $A_f$ | $k$ | $\tau_0$ |
| --- | --- | --- | --- |
| eve2:neutral | $0.44 \pm 03$ | $11.8 \pm 1.3$ | ↑ |
| eve2:wt | $0.90 \pm 03$ | $4.0 \pm 0.3$ | |
| eve2[Zld]:neutral | $0.86 \pm 03$ | $4.3 \pm 0.5$ | $210 \pm 20 \text{ s}$ |
| eve2[Bcd]:neutral | $0.74 \pm 04$ | $8.2 \pm 1.1$ | ↓ |
| eve2[Dst]:neutral | $0.93 \pm 02$ | $8.2 \pm 0.9$ | |

From the model, the first passage distributions can be explained by a varying number of rate limiting steps, each with a characteristic time of 210 ± 20 s. Each of the activating TF reporters increased the active fraction of nuclei within the stripe relative to *eve2:neutral* (in Table 1). This is consistent with what would intuitively be expected from inspection of the dynamic transcription profiles in Fig. 1D. In addition, all reporters decreased the number of rate limiting steps relative to *eve2:neutral*: = 11.8 ± 1.3 for *eve2:neutral*, 8.2 ± 1.1 and 8.2 ± 0.9 for *eve2[Bcd]:neutral* and *eve2[Dst]:neutral*, respectively, and 4.0 ± 0.3 and 4.3 ± 0.5 for *eve2:wt* and *eve2[Zld]:neutral*, respectively. Regardless of the detailed interpretation of this model, it is clear that while all reporters reduce the number of rate limiting steps on the path to first passage transcription activation, those that bind Zld (*eve2:wt* and *eve2[Zld]:neutral*) do so greatly. To a large extent, this explains the differences in the dynamic transcription profiles of Fig. 1D, but not entirely. The activating TF reporters must be acting to tune other kinetic parameters in addition to the first passage time.

#### Active transcription lifetimes

The lifetime of a single active transcription interval is the time over which transcription signal is continuously detected within a single nucleus (Fig. 3A); this value is proportional to the number of RNA molecules synthesized during that interval (Bothma et al., 2014). The cumulative distributions in Fig. 3B show the frequency and survival of the active transcription lifetimes.

These curves can reveal one or more *types* of activations present in the distribution, each type defined by a characteristic lifetime (Friedman and Gelles, 2015). For example, the distributions in Fig. 3B do not reflect the presence of a single characteristic lifetime (SFig. 4). A bi-exponential probability density function, which is commonly used to describe chemical kinetics (e.g. (Friedman and Gelles, 2012)), was well suited to model these distributions:

**Figure 4.**
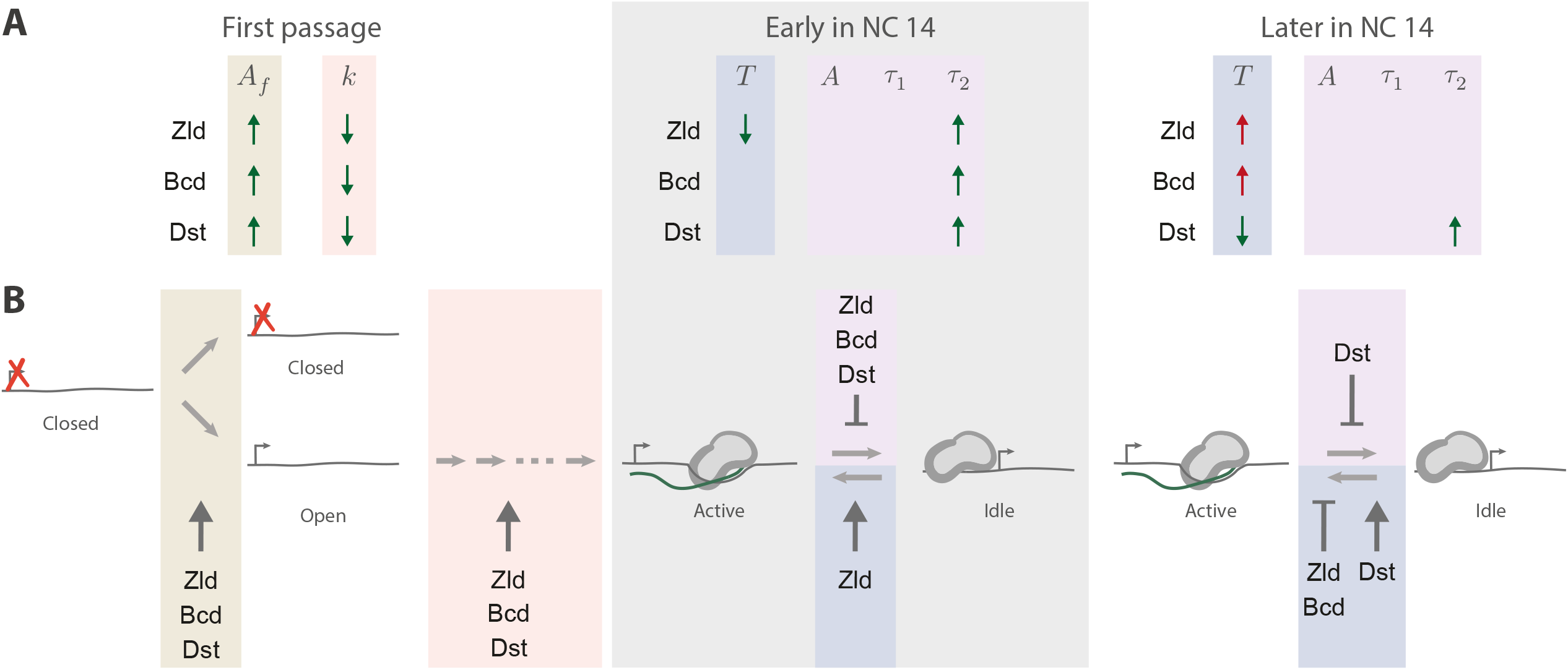
Kinetic roles of three transcriptional activators. (A) The effect of each activating TF on each kinetic parameter. The arrows denote if a parameter is increased or decreased by a TF. Parameter changes consistent with activation are shown in green and changes consistent with repression in red. Beige and peach mark the first passage transcription model parameters (Eq. 1), blue marks the idle transcription model parameter (Eq. 3), and purple the active transcription model parameters (Eq. 2). The gray shaded region denotes active and idle transcription parameters early in NC 14. (B) Model of transcription and its regulation during NC 14. Colored regions are as in (A). Dark arrows indicate an activator-induced decrease in the average idle period (blue), decrease in the number of rate limiting steps (peach), or an increase in the active fraction of nuclei (beige). T-bars indicate an activator-induced increase in the characteristic active transcription lifetime (purple) and average idle period (blue). The ellipsis represents a variable number of rate limiting steps on the pathway to first passage transcription.

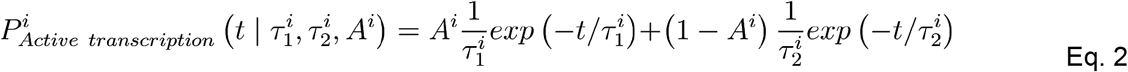

Where 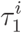 and 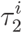 are characteristic lifetimes, *A*^*i*^ and 1 − *A*^*i*^ are the relative amplitudes of each, respectively (Fig. 3B, smooth curves). The parameters 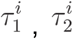 and, *A*^*i*^(Table 2) were determined by maximum likelihood fitting to the distributions of active transcription times (see Methods). The two exponential terms reflect two types of active transcription lifetimes by both *eve2:neutral* and *eve2:wt*: a short-lived type characterized by 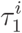 and a long-lived type characterized by 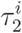. Thus, once activated, transcription does not turn off stochastically with a single characteristic lifetime. Kinetically, each of these activation types represents a kinetic state of the system, each of which contribute to the two terms in Eq. 2. The exact details of the kinetic stability, the pathways into and out of, and the biochemical identities of these states cannot be determined from the data and analysis here.

**Table 2.** Model parameters values for active transcription. See Eq. 2. Early lifetime distributions are composed of the first 20% of active transcription intervals detected for each reporter. Later distributions are composed of the other 80%. Standard errors were computed using bootstrapping methods.

| | | $A$ | $\tau_1$ | $\tau_2$ |
| --- | --- | --- | --- | --- |
| All lifetimes | eve2:neutral | $0.53 \pm 0.23$ | $54 \pm 26$ s | $190 \pm 60$ s |
| | eve2:wt | $0.60 \pm 0.07$ | $71 \pm 15$ s | $460 \pm 60$ s |
| | eve2[Zld]:neutral | $0.54 \pm 0.18$ | $61 \pm 30$ s | $260 \pm 70$ s |
| | eve2[Bcd]:neutral | $0.55 \pm 0.11$ | $64 \pm 16$ s | $280 \pm 40$ s |
| | eve2[Dst]:neutral | $0.55 \pm 0.06$ | $61 \pm 15$ s | $560 \pm 60$ s |
| Early lifetimes | eve2:neutral | $0 \pm 0.18$ | N/A | $148 \pm 51$ s |
| | eve2:wt | $0.30 \pm 0.10$ | $45 \pm 21$ s | $530 \pm 80$ s |
| | eve2[Zld]:neutral | $0.34 \pm 0.34$ | $136 \pm 100$ s | $370 \pm 80$ s |
| | eve2[Bcd]:neutral | $0.37 \pm 0.22$ | $44 \pm 24$ s | $360 \pm 140$ s |
| | eve2[Dst]:neutral | $0.32 \pm 0.10$ | $40 \pm 17$ s | $630 \pm 120$ s |
| Later lifetimes | eve2:neutral | $0.49 \pm 0.17$ | $50 \pm 36$ s | $160 \pm 20$ s |
| | eve2:wt | $0.54 \pm 0.09$ | $65 \pm 43$ s | $290 \pm 20$ s |
| | eve2[Zld]:neutral | $0.36 \pm 0.33$ | $39 \pm 28$ s | $140 \pm 40$ s |
| | eve2[Bcd]:neutral | $0.42 \pm 0.23$ | $51 \pm 32$ s | $190 \pm 60$ s |
| | eve2[Dst]:neutral | $0.48 \pm 0.10$ | $50 \pm 27$ s | $380 \pm 40$ s |

The model in Eq. 2 reveals the kinetic differences between activation by *eve2:wt* and activation by *eve2:neutral*. The relative amplitudes of the short-lived active transcription lifetimes, *A* ^*eve2:wt*^ and *A*^*eve2:netural*^, are equivalent within error. This suggests that in each case, the distribution is composed of an equivalent fraction of short-lived and long-lived active lifetimes. The apparent increase in active lifetimes driven by *eve2:wt* is due to an increased characteristic lifetime for the long-lived activations by *eve2:wt* (e.g.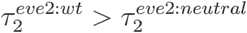, Table 2).

For both *eve2:wt* and *eve2:neutral* the biexponential nature of the active transcription lifetime distributions can be, at least in part, attributed to a time-dependent characteristic lifetime, with longer activations more likely to occur early in the nuclear cycle and shorter activations more likely to occur later. We divided the lifetime measurements of Fig. 3B into two subsets: those that first activated early in NC 14 and those that first activated later (Fig. 3C and SFig. 5, respectively; see Methods). We applied the model in Eq. 2 to each of the two subset distributions (early and later) of *eve2:wt* and *eve2:neutral*. These subsets yielded characteristic lifetimes, 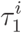 and 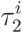, equivalent to those of the whole distributions (Table 2). However, the relative amplitudes, *A*^*i*^ and 1 − *A*^*i*^, were different. In each case the relative amplitude for the early distribution was dominated by long-lived activations (i.e. 1 − > *A*^*i*^). For *eve2:wt*, the amplitude of long-lived activations for the early distribution was, 1 − *A* ^*eve2:wt, early*^=0.70 ±0.80, up from 1 − *A* ^*eve2:wt*,^ =0.40 ±0.07 for the whole distribution. For *eve2:neutral* the long-lived amplitude was 1 − *A* ^*eve2:netural,early*^ =1.00 ±0.18 for the early subset, compared to 1 − *A* ^*eve2:netural*^ =0.53 ± 0.23 for the whole distribution. However, the biexponential nature of these distributions may not be entirely attributable to the time-dependency of the characteristic lifetimes, which we address in the Discussion below.

We determined how additional activity of Zld, Bcd and Dstat in the activating TF reporters affected the characteristic active lifetimes by applying the model of Eq. 2 to their respective distributions (Fig. 3B). This revealed that, for the undivided distributions, only *eve2[Dst]:neutral* displayed active lifetime kinetics that were different from *eve2:neutral* (Table 2). *eve2[Dst]:neutral* induced longer long-lived active times 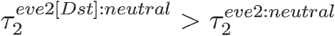. Upon further analysis, when these active lifetimes were divided into early and later subsets, this trend held up for the later active lifetime distributions (SFig. 5, Table 2). However, for the subset of early active times, all three activator reporters showed an increase in the characteristic lifetime for long-lived active times, 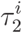 (Fig. 3C, Table 2). From this, two conclusions can be drawn. First, the activator reporters increase active lifetimes by altering the long-lived characteristic time, 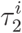, and not the short-lived characteristic time,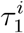, nor the fraction of long-lived events, 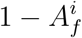. Second, the differences in active lifetimes predict that at any given moment, more nuclei will be undergoing active transcription for each of the activating TF reporters when compared to *eve2:neutral*. This further explains the differences in the dynamic transcription profiles of Fig. 1D.

#### Idle transcription

We define idle transcription as the interval of time between the end of one observed active transcription interval and the beginning of the next within the same nucleus (Fig. 3D, SFig. 6). During these intervals, the MCP-GFP transcription signal was below detection threshold. Analogous to a car engine that idles while waiting at a stop light, transcription may still be occurring at a low level during these periods, but at a level that we could not detect. As with active transcription lifetimes, idle periods are related to the number of transcripts synthesized over NC 14; decreasing the mean idle period increases the total time over which a locus is actively transcribing. Moderating the length of idle periods in a time-dependent fashion can regulate when and how much of a transcript is produced.

The inverse of the mean idle period, with units of s^-1^, can be read directly from the cumulative frequency distributions in Figs. 3B, 3C, and SFig. 5 via the horizontal axis intercept. This value is calculated by:

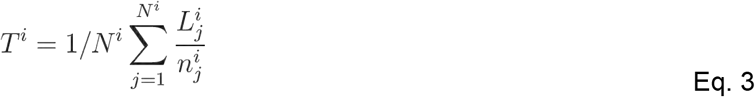

Where is *N*^*i*^ the total number of nuclei with at least one detected active transcription interval, *L*^*i*^ is the cumulative length of time during which no transcription signal was detected in these same nuclei (see SFig. 6), and *n*^*i*^ is the number of active transcription intervals (of any length) observed in these nuclei.

The characteristic idle period is an order of magnitude shorter early in NC 14 than it is later for all reporters (Fig. 3E and F; Table 3). Compared to *eve2:neutral, eve2[Zld]:neutral* shortens the idle period early in NC14, while *eve2[Dst]:neutral* does so later. It might be expected that the active transcription times and the idle periods are regulated in concert: as the former increases, the latter decreases. This is not the case. For example, later in the nuclear cycle *eve2[Zld]:neutral* and *eve2[Bcd]:neutral* drive longer idle times than *eve2:neutral* (Table 3) while the active transcription parameters are identical within error during that time (Table 2). In these data, the idle period is tuned independently of active transcription lifetimes and varies over the nuclear cycle, with dramatically shorter periods early in NC 14.

**Table 3.** Mean idle transcription periods. See Eq. 3. Early idle times are the first 20% detected for each reporter. Later idle times are the other 80%. Standard errors were computed using bootstrapping methods.

| | | $T$ |
| --- | --- | --- |
| All idle times | eve2:neutral | $280 \pm 30$ s |
| | eve2:wt | $260 \pm 10$ s |
| | eve2[Zld]:neutral | $410 \pm 30$ s |
| | eve2[Bcd]:neutral | $400 \pm 30$ s |
| | eve2[Dst]:neutral | $230 \pm 10$ s |
| Early idle times | eve2:neutral | $59 \pm 14$ s |
| | eve2:wt | $46 \pm 5$ s |
| | eve2[Zld]:neutral | $20 \pm 3$ s |
| | eve2[Bcd]:neutral | $37 \pm 6$ s |
| | eve2[Dst]:neutral | $48 \pm 6$ s |
| Later idle times | eve2:neutral | $280 \pm 30$ s |
| | eve2:wt | $240 \pm 10$ s |
| | eve2[Zld]:neutral | $410 \pm 30$ s |
| | eve2[Bcd]:neutral | $370 \pm 30$ s |
| | eve2[Dst]:neutral | $200 \pm 10$ s |

## DISCUSSION

Our goal was to determine the kinetic roles of three different TFs known to activate transcription. We measured their effects on transcription in living *Drosophila blastoderm* embryos using multiple permutations of an *eve2* reporter driving MCP-GFP marked nascent transcripts, and models derived from chemical kinetics. We characterized two benchmark reporters that served as effective lower- and upper-bounds, which allowed us to compare their output to reporters that encode additional binding motifs for specific activator proteins. Straightforward inspection of the data yielded qualitative insights into roles for the three TFs: Zld, Bcd and Dstat. Quantitative analysis with our empirical models provided additional insight by contrasting the probability distributions of first passage activation times, active transcription lifetimes, and idle transcription periods associated with each transgenic reporter. We found that the active transcription distribution and mean idle period change over time, with longer activation lifetimes and shorter idle periods early in nuclear cycle 14. We further found that each TF drove transcription in a kinetically unique way, as laid out in Fig. 4 and discussed in more detail below. Our results also highlight unresolved questions of eukaryotic gene regulation, including how TFs bound outside of enhancers affect transcriptional output and how different combinations of TFs can achieve similar transcription output.

### Insights into transcriptional kinetics

The models used here are agnostic to the underlying biochemical states of the transcriptional system. In effect, we are summarizing the kinetics of transcription *signals* by using empirical models to describe probability distributions. The time constants reported here are related, but not identical, to the underlying kinetic pathway of transcription. This is in contrast to recent work that assumed an underlying model of transcription then measured transitions between states by inferring the state of the system from the intensity record of a fluorescent MS2/MCP reporter (Lammers et al., 2020). In this instance, we sought to quantify how transcriptional outputs changed with increased Zld, Bcd, and Dstat activity. Our approach is mathematically straightforward, with well established methods for its application (Friedman and Gelles, 2015; Zhou and Zhuang, 2007), and it relies on a simple binarization of MS2/MCP data. It is broadly applicable to all live imaging transcription studies.

Our analysis was restricted to the distributions of first passage activation times, active transcription lifetimes, and mean idle period. Other distributions, including fluorescence intensity (signal brightness) and intensity fluctuations (noise), were not included due to the dependency of these measurements on experimental conditions that are difficult to control. These include excitation laser drift and day-to-day variability, heterogeneous illumination across the field of view of the microscope, nucleus-to-nucleus variation in the depth of the reporter gene relative to the surface of the embryo, and photobleaching of GFP fluorophore over the course of an experiment. All of these sources of extrinsic noise skew fluorescence intensity measurements and must be accounted for when reporting those distributions. These variables can and should be controlled for to the extent that they can be. However, because we largely eschew measures of signal intensity, the approach laid out here is robust against them.

We believe this is the first report of a bi-exponential distribution of active lifetime MS2/MCP measurements. A bi-exponential probability distribution implies that multiple stable states of the system exist, each with a discernible characteristic lifetime (Friedman and Gelles, 2015). These distributions change over time (Fig. 3B, 3C, and SFig. 5). One possible explanation is that transcription kinetics change as extrinsic inputs change over the course of NC 14. For example, nuclear TF concentrations are not static over the nuclear cycle (Bothma et al., 2018). It is also possible that the bi-exponential nature is attributable to an unknown artifact in either the MS2/MCP measurements or the detection methods used here. If that were to be the case, our conclusions likely would not change. Significant differences between active lifetime distributions were almost entirely attributable to changes in the long-lived characteristic lifetime,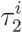. Unless some experimental artifact systematically altered these stable, long-lived transcription signals, we would have reached the same conclusions.

One striking result from our study is the large differences in the time delays between the end of anaphase and the first detection of transcription. These differences strongly suggest that the regulation of the rates of first passage transcription is an important and widespread mechanism of activating TFs. From qualitative inspection, each of the activating TFs reduced this delay, and they did so in dramatically different fashions (Fig. 1D). Our modeling approach expounded on this conclusion, and found that each TF not only increased the active fraction of nuclei, which had previously been shown to be important for patterning in the embryo (Garcia et al., 2013; Lammers et al., 2020), but also decreased the number of rate limiting steps,. From this, each TF is implicated in regulating the kinetic steps that lead to the onset of transcription, and not just the likelihood that a locus ever turns on. It may be that this is a role played by all activating TFs. These findings are a strong argument for pairing mathematical modeling with dynamic imaging and provide an exciting prospect for further study to determine the importance and extent of first passage regulation by transcription factors.

### Insights into TF function

Zld is thought to act as a pioneer factor in the embryo, opening chromatin and maintaining it in a state competent for transcription (Harrison et al., 2011; Schulz et al., 2015; Sun et al., 2015). Kinetically, it is reasonable to hypothesize that this mechanism would decrease the number of rate limiting steps to first passage transcription. Consistent with this, we found that Zld dramatically decreased the number steps, *k*^*i*^. However, Zld also affected other kinetic parameters (Fig. 4). Zld increased transcription by increasing active transcription lifetime, *τ*_2_, and decreasing the idle period, *T*, but only for early times in the nuclear cycle. Zld has been shown to perform multiple roles at different target genes (Dufourt et al., 2018; Eck et al., 2020), but to our knowledge, this is the first evidence that it plays multiple roles at a single gene.

Two Zld binding site insertions served to turn the dynamic transcription profile of *eve2:neutral* into something close to that of *eve2:wt* (Fig. 1D). From this qualitative inspection of the data, one might conclude that the similarity is due to a similar number of Zld binding motifs in both *eve2[Zld]:neutral* and *eve2:wt* (see the sequence schematics in Fig. 1A), even though regulatory sequences are sensitive to binding motif position, orientation, and neighboring sequences (Erceg et al., 2014; Vincent et al., 2016). However, our modeling does not support this naive interpretation. Following first passage into active transcription, *eve2:wt* maintained active transcription by increasing the active transcription lifetimes, *τ*_2_, over all of NC 14. Zld did so by increasing *and* decreasing the idle period, *T*, but only for early times in the nuclear cycle. Zld then repressed active transcription later on by increasing the idle period. Therefore, the similarity between the *eve2[Zld]:neutral* and *eve2:wt* dynamic transcription profiles cannot entirely be attributed to Zld activity as the two regulatory sequences produce similar spatiotemporal outputs by acting on different combinations of kinetic parameters.

Bcd is a highly studied TF, with considerable focus placed on how the Bcd gradient and cooperative interactions between Bcd proteins regulate target genes (Burz et al., 1998; Driever and Nüsslein-Volhard, 1988; Hannon et al., 2017; Lebrecht et al., 2005; Park et al., 2019; Struhl et al., 1989). Our characterization of kinetic roles found that Bcd, in this context, increased the active fraction of nuclei and decreased the number of rate limiting steps to first passage transcription. This is, perhaps, consistent with a previous report of Bcd activity early in the nuclear cycle and its capacity for pioneering activity (Hannon et al., 2017). However, later in the nuclear cycle, Bcd suppressed transcription by increasing the idle transcription period (Fig. 4). This may be unsurprising, given another previous report that Bcd is a bifunctional activator in certain contexts (Liaw and Lengyel, 1993), but the activating/suppressing activities of Bcd reported here are different than bifunctional regulation. In this context, Bcd acts to tune different parameters at different times during the nuclear cycle, sometimes resulting in an increase in the number of transcripts synthesized, and other times resulting in a decrease. Although intriguing, these conclusions should be treated with caution. In this context, the effect of additional Bcd motifs was relatively small. Of the three activating TF reporters, the dynamic transcription profile of *eve2[Bcd]:neutral* was most similar to the baseline profile of *eve2:neutral* (Fig. 1D). This was to be expected for two reasons. First, Bcd has been characterized as a weak activator (Ma et al., 1996, 1999). Second, of the three TFs tested here, *eve2* contains the most native binding motif sequences for Bcd (Fig. 1A). Here, Bcd activity may be close to saturation. An alternative variant of the *eve2* reporter with fewer Bcd binding sites may test this hypothesis and give further insight into the kinetic role of bicoid.

Dstat is ubiquitously expressed in the embryo, is known to activate *even-skipped* stripe 3 and 5 (Fujioka et al., 1999; Struffi et al., 2011; Yan et al., 1996), and is thought to activate all *even-skipped* enhancers (Barr and Reinitz, 2017). There are two predicted binding motifs in *eve2* and four, albeit weaker, predicted sites in the endogenous spacer sequence (Fig. 1A). Given this, it was somewhat surprising that *eve2[Dst]:neutral* drove a dynamic transcription profile that differed starkly from both *eve2:neutral* and *eve2:wt*. Dstat acted on nearly every kinetic parameter in our models (Fig. 4). These results establish a role for Dstat during initial activation of a locus: increasing the active fraction of nuclei and decreasing the number of rate limiting steps to first passage transcription. Additionally, unlike both Bcd and Zld, Dstat activated transcription throughout the nuclear cycle by both increasing active lifetimes and decreasing the idle periods between those lifetimes.

## Conclusion

A mandate of systems biology with respect to transcriptional regulation is to decipher the logic of transcriptional control and predict regulatory sequence function (Catarino and Stark, 2018). This requires establishing why specific TFs have been selected to operate at a particular gene at a particular time. How is the means by which a TF induces transcription consequential? To answer this the mechanisms of activation must first be established. TF mechanisms have always been defined by the methods with which we characterize them. Genetic approaches establish proteins as activating or repressing, biochemistry identifies the complexes a protein interacts with, and genomics establishes the genetic targets of a protein. Each approach plays a role in unveiling the mechanisms by which TFs regulate transcription. A niche of live imaging —by MS2/MCP and other methods—is to define *kinetic* mechanisms of regulation. The shortcoming of this approach is its scalability. It is difficult to imagine applying this approach to, for example, the hundreds of human TFs (Lambert et al., 2018). This is, however, a tractable task within the blastoderm with its ∼40 TFs present (MacArthur et al., 2009), once again placing the blastoderm as a model for higher organisms (Gregor et al., 2014).

The results presented here complicate conventional mechanistic labels for TFs such as activator, repressor, pioneer factor, bifunctional factor. In this instance, Bcd and Zld act to unequivocally increase transcriptional outputs, but do so by both activating some kinetic steps while repressing others (Fig. 4). Does this make them activators or bifunctional factors? Each TF plays a significant role during first passage activation, does this qualify each of them as a pioneer factor? Zld has a complicated kinetic role throughout the nuclear cycle that includes more than what might reasonably be attributed to a pioneer factor. Defining TFs by their kinetic roles skirts these ambiguities. More importantly, it builds upon a foundation with which we can work towards predicting transcriptional outputs *a priori*.

## METHODS

### Cloning and transgenesis

An MS2 transcription reporter gene was placed in the pBOY vector backbone (Hare et al., 2008) using Gibson isothermal assembly (Gibson et al., 2009). The reporter consisted of the *Drosophila melanogaster even-skipped* core promoter, a 1295 bp sequence encoding 24 tandem repeats of MS2 stem loops ((Hocine et al., 2013); Addgene #45162), 3 kb of the lacZ gene, and the alpha-tubulin 3′ UTR. For *eve2:neutral* (Addgene accession number pending), we computationally designed a sequence predicted to lack binding motifs for regulatory proteins active in patterning the blastoderm embryo, motifs for architectural binding proteins, and core promoter sequences using the online binding motif removal tool SiteOut (Estrada et al., 2016; Scholes et al., 2019). This sequence, along with the 484 basepair minimal *eve2* sequence (Small et al., 1992), was commercially synthesized (GenScript gene synthesis services) and cloned into our reporter plasmid through isothermal assembly (Gibson et al., 2009). For *eve2:wt* (Addgene number pending), the 1033 bp that separate minimal *eve2* and the *even-skipped* promoter were PCR amplified from genomic DNA, then inserted into our reporter plasmid along with *eve2* also using isothermal assembly. For *eve2[Zld]:neutral* (Addgene number pending), a sequence identical to *eve2* save for two motif mutations—tccgccgat became tAATccgat at 299 bp with respect to the 5′ end of *eve2* and ttctgcggg became ttAATcCgg at 323 bp—was synthesized and cloned as above. For *eve2[Bcd]:neutral* (Addgene number pending), the mutations to *eve2* were: tccgccgat became tcAgGTATt and ttctgcggg became ttcAgGTAg. For *eve2[Dst]:neutral* (Addgene number pending) the mutations to *eve2* were: tccgccgat became tTcCcGgaA and ttctgcggg became ttcCCGgAA.

We verified the sequence of the enhancers and promoter of all reporter constructs prior to injection, and checked the length of the MS2 cassette by restriction digest. The pBOY backbone contains an attB site for phiC31-mediated site-specific recombination (Fish et al., 2007) and a mini-white gene for transformant selection. For each construct, BestGene Inc. (Chino Hills, CA) injected midi-prepped DNA into 200 embryos of Bloomington Stock BL8622, which contains the attP2 landing site on chromosome 3L (Markstein et al., 2008). All constructs are integrated into this same attP2 landing site. After the constructs were successfully integrated into the fly genome, we prepared genomic DNA, PCR-amplified the transgene and repeated the sequencing and restriction digest verification of the reporters.

### Live imaging

Virgin females with the genotype yw; His2Av-mRFP1; MCP-NoNLS-eGFP (Garcia et al., 2013) were crossed to males homozygous for one of the transgenic transcription reporters. Embryos no older than 30 minutes were collected and subsequently dechorionated in freshly-made 50% bleach for two minutes. Embryos were placed on a single coverslip and bathed in Schneider’s *Drosophila* medium (Gibco), where they remained for the entire duration of the experiment.

Light sheet microscopy was performed on a diSPIM (Applied Scientific Instrumentation, Eugene OR) setup as previously described (Kumar et al., 2014), though only a single imaging view was used for all experiments presented here. 488 & 561nm laser lines from an Agilent laser launch were fiber-coupled into MEMS-mirror scanhead, used to create a virutally-swept light sheet. A pair of perpendicular water-dipping, long-working distance objectives (NIR APO 40×, 0.8 NA, Cat. No. MRD07420; Nikon, Melville, NY) were used to illuminate the sample and to collect the resulting fluorescence. All laser lines were reflected with a quad-pass ZT405/488/561/640rpcv2 dichroic and emission was selected with a ZET405/488/561/635M filter (Chroma) before detection on a sCMOS camera (ORCA Flash v2.0; Hamamatsu). For data acquisition and instrument control, we used the ASI diSPIM plugin within MicroManager (Edelstein et al., 2014).

Image acquisition commenced during NC 13 and ceased at the beginning of gastrulation. Z-stacks were acquired every 30 seconds by sweeping the sheet in conjunction with the detection plane (controlled via piezo motor) through the sample. Z-stacks were composed of 80 Z-planes separated by 0.5 μm; the exposure time to collect a single Z-plane was 50 milliseconds. Two embryos were imaged for each transgenic reporter.

### Image analysis

Image analysis was done using custom software implemented in Matlab. Algorithms for automatic spot and nuclei detection and tracking were adapted from (Friedman and Gelles, 2015). Following maximum intensity projection of mRFP1 and eGFP emission Z-stacks for each time frame, the nuclei were segmented. Spots of transcription were located in each time frame using an automated spot detection algorithm that considered spot intensity, shape, and hysteresis (Friedman and Gelles, 2015). Each spot was fit with a 2D gaussian to determine the center coordinates then associated with the closest nucleus. Cases where multiple spots were associated with the same nucleus in the same frame were rare (< 10 instances per data set). These were resolved by hysteresis: the spot closest to the location of spots associated with that same nucleus in adjacent time frames was chosen. A nucleus was considered actively transcribing at a given time frame if a transcription spot of GFP fluorescence was detected within it.

### Determining model parameters

All modeling was restricted to nuclei located in the center of the stripe 2 domain. To determine those nuclei, the time-dependent mean position of each nuclei was computed in units of percent of anterior-posterior axis length (AP). This mean position was computed over the time interval starting with the appearance of the first active transcription spot and ending with the disappearance of the last active spot within a single embryo. Every nucleus with a mean position within 2% AP of the mid-point of the stripe was considered within the center of the stripe and was included in the modeling analysis (SFig 1).

To derive the model parameters of Eq. 1, we used maximum likelihood methods to directly fit the underlying active transcription observations as described in (Friedman and Gelles, 2015). This method accounts for limited resolution due to digital representation of real numbers and finite observation times. We did not directly fit the cumulative frequency distribution using conventional fitting procedures that assume independent errors because each point in the curve includes the random errors of all points to the left. The data from two imaging replicates for each transgenic reporter were combined and treated as a single dataset, as is typical for analysis of fluorescence spectroscopy experiments (e.g. (Harden et al., 2016)).

The mean idle period values (Eq. 2) were computed for each transgenic reporter by summing the total time that transcription was not detected within each active nucleus. This total inactive time included both the time between active transcription intervals as well as the time between the end of the final active interval and gastrulation (SFig. 6, black regions). The total inactive time,*L*^*i*^, was then divided by the number of inactive intervals, *n*^*i*^.

To determine the first passage model parameters (Eq. 3), we again used the maximum likelihood methods described in Friedman et al. 2015. We jointly fit the gamma scale parameter, τ_0_, to all conditions while simultaneously fitting the active fraction, 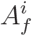, and the gamma shape parameter,*k*^*i*^, to each condition individually. We maximized the likelihood function:

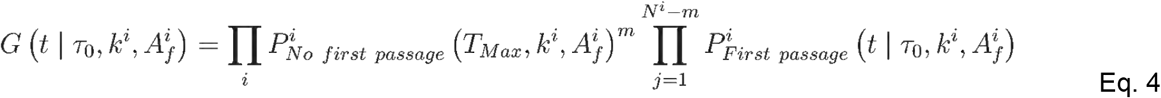

Where *i* is each condition *eve2:wt, eve2:neutral, eve2[Zld]:neutral, eve2[Bcd]:neutral*, and *eve2[Dst]:neutral*.*T*_*Max*_ is the maximum observation time, i.e. the length of NC 14. *N*^*i*^ is the total number of nuclei within the center of the stripe.*m* is the number of nuclei located in the center of the stripe but in which a transcription spot never appears.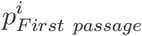 is given in Eq. 3.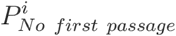 is the probability a nucleus is active but does not display a transcription spot during NC 14. This accounts for the stochastic reality that, under a given set of kinetic parameters, it is possible that a nucleus does not display transcription because it does not have time to turn on before gastrulation. In this possibility, a nucleus is not being actively suppressed nor does it lack sufficient activating TF activity. Thus we consider it active, despite a lack of transcription signal. This term is necessary to accurately determine the active fraction of nuclei,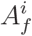.Because of the limited precision of digital data, we maximized the sum of the logarithms of the individual factors of Eq. 4 instead of their product (Friedman and Gelles, 2015).

To make the “early” distribution of active transcription lifetimes (Fig. 3C, Table 2), we selected the 20% of lifetimes that first appear in any nucleus for each imaging replicate (e.g. 61/307 lifetimes for one replicate of *eve2:wt*). The remaining active transcription observations made up the “later” distribution (SFig. 5, Table 2). The same method was used to separate the idle period distributions into early (Fig. 3F, Table 3) and later (Table 3) subset distributions.

### Error analysis

All kinetic parameter error distributions were estimated by bootstrapping (Efron and Tibshirani, 1994). Briefly, for each measurement (e.g. active transcription lifetimes, idle periods, first passage time) we generate 10,000 simulated data sets for each genetic condition. To generate these, we randomly sample with replacement from the experimental observations. Bootstrapping of the first passage time distribution is an illustrative example. The *eve2:wt* experimental data set contained 88 nuclei within the center of the stripe. From these 88 nuclei, 88 observations were made. Some of these observations were a first passage time (from nuclei that displayed at least one instance of active transcription) and some of these observations represented nuclei that never displayed transcription. Thus each simulated *eve2:wt* data set contained 88 observations drawn from the experimental one. These simulated data sets were subsequently analyzed with the exact methods that were applied to the experimental data sets, as described above. A distribution of values is thus generated for each kinetic parameter. Standard statistical methods were then used to find the standard deviation of each parameter. We report these as error values in all tables and the bar charts in Fig. 3.

The shaded error regions of the frequency survival plots in Fig. 3B, 3C, and SFig 5 and the cumulative first passage plot in Fig. 2 were also determined by bootstrapping. These regions represent the 90% confidence intervals, i.e. 90% of the simulated datasets fall within this range.

## ACKNOWLEDGEMENTS

We thank Tally Lambert and Jennifer Waters at The Nikon Imaging Center at Harvard Medical School for guiding the microscopy experiments, assistance in data processing, and lively discussions. We also thank Larry Friedman, Jane Kondev, members of the DePace lab, and our reviewers for helpful comments and feedback.

## AUTHOR CONTRIBUTIONS

All authors designed the research. T.H. and B.V. performed the research. T.H. analyzed the data and drafted the manuscript. All authors contributed to writing the final version of the manuscript.

---

**Supplemental Figure 1.**
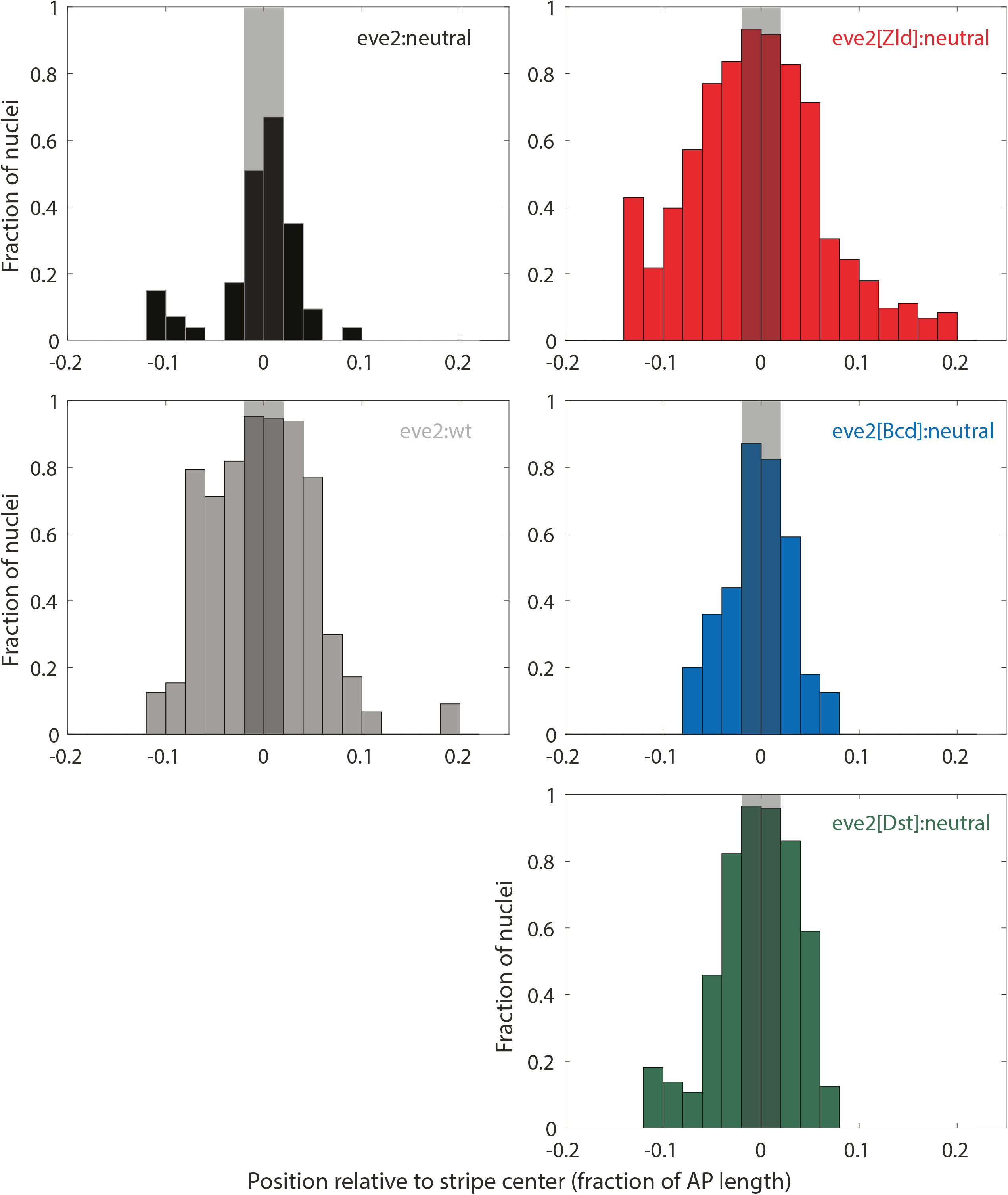
Kinetic analysis was restricted to nuclei located in the center of stripe 2. Histograms of the fraction of active nuclei in a given spatial region of the embryo. Active is defined as having at least one instance of active transcription over the course of NC 14. Nuclei were binned according to their mean location over NC 14, shown on the horizontal axis in units of percent of the anterior-posterior axis length (see Methods). The center of the stripe is located at 0 on the horizontal axis. Gray shaded regions show the location of the nuclei analyzed in Figs. 2 and 3. Repressors, including Giant and Kruppel, act around either edge of the stripe to set the boundaries. Thus, analysis excluded these edge regions in an attempt to isolate activating TF activity from repressive TF activity.

**Supplemental Figure 2.**
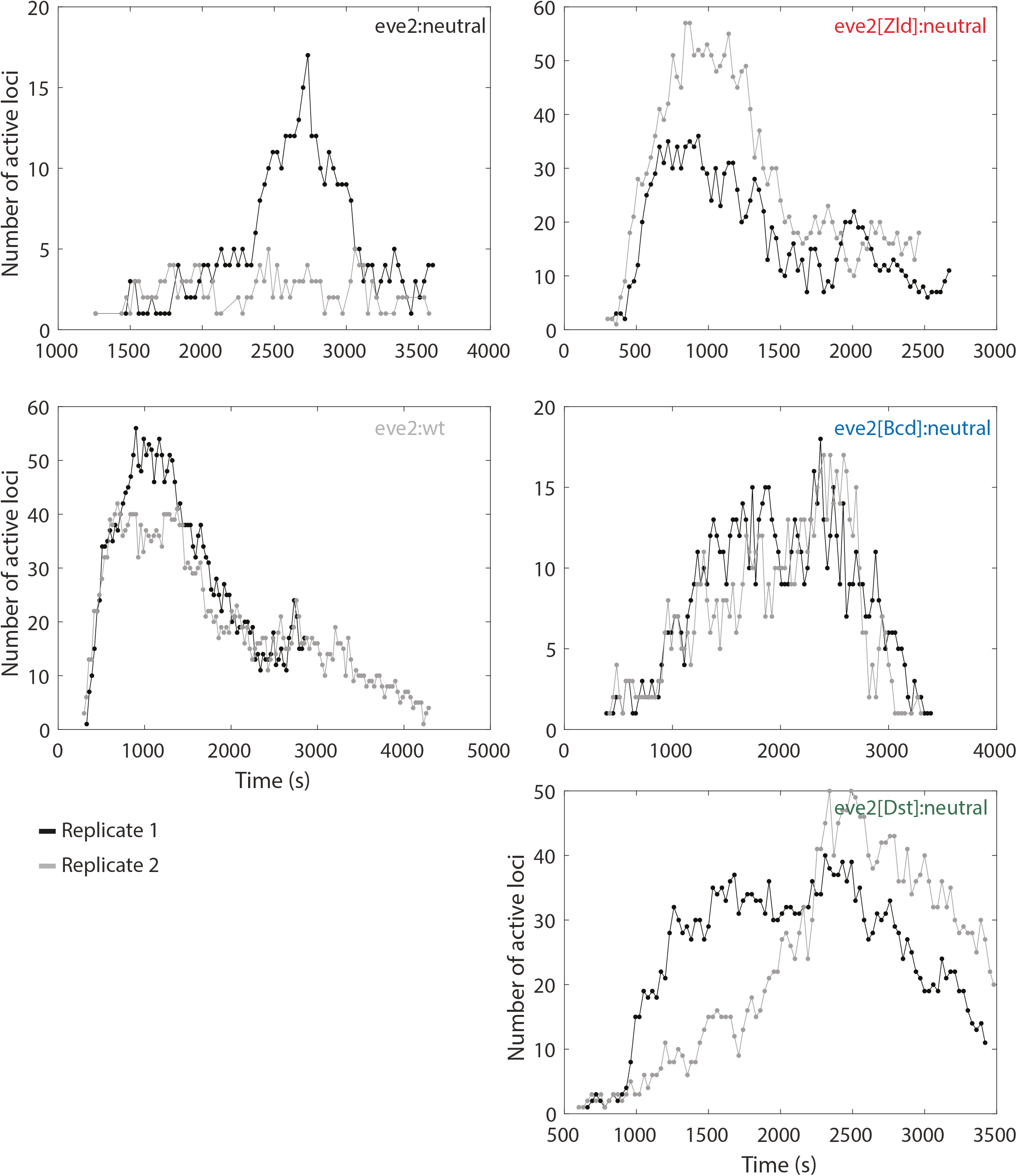
Reproducibility of the dynamic transcription profile between biological replicates. The dynamic transcription profiles, as in Fig. 1D, for two different replicates of each transcription reporter (gray and black).

**Supplemental Figure 3.**
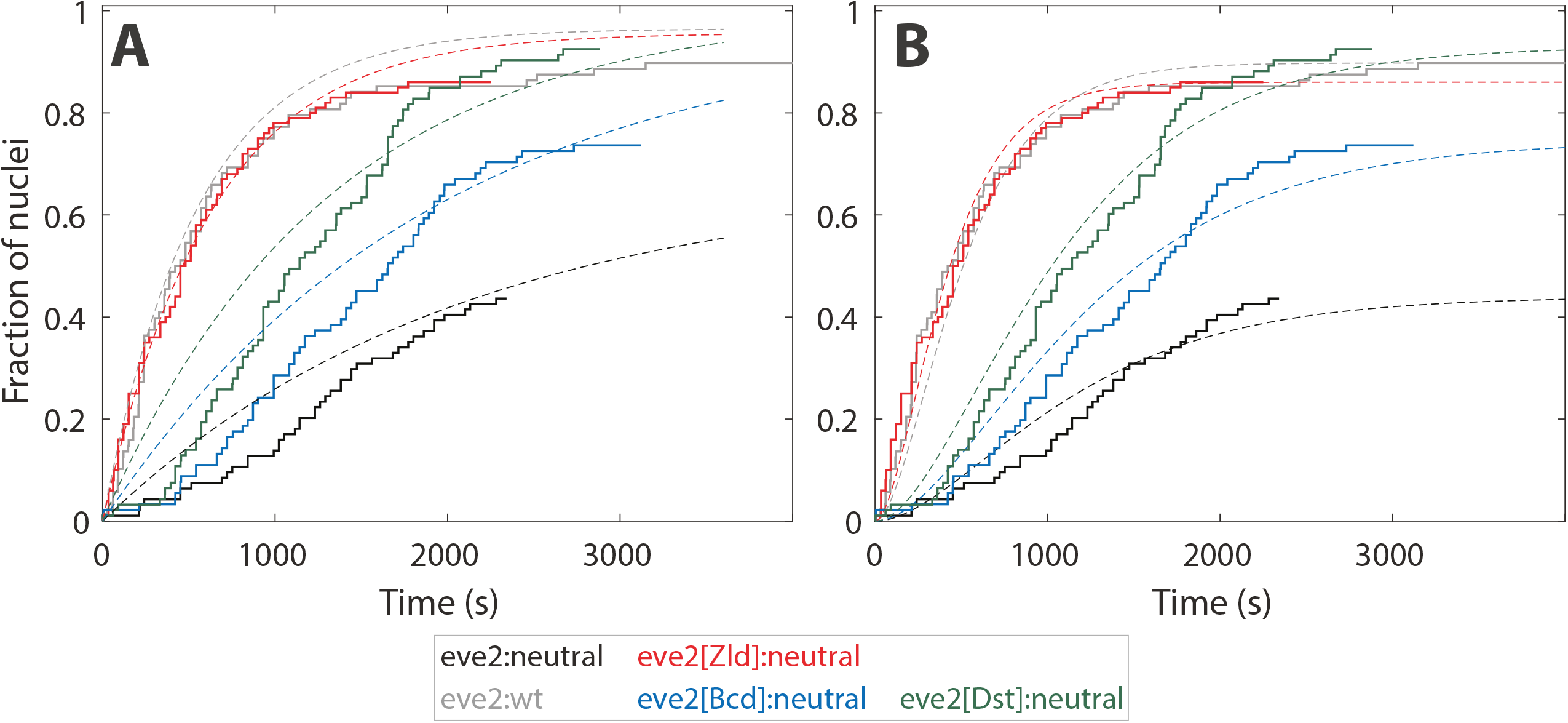
Chemical kinetics models typically used to measure first passage rate constants cannot describe first passage into active transcription. (A) Cumulative first passage distributions (solid lines) as in Fig. 2B, but here the initial time delay between the end of anaphase and the first detection of transcription is ignored; *t* = 0 is the time at which a transcription signal was first detected in the embryo, as has been done previously (Dufourt et al., 2018). The distributions are overlaid with a single step association model:

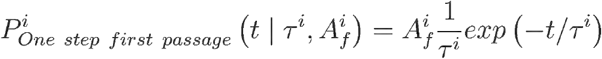

with an active fraction and characteristic time fit parameter,τ and 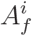, respectively (dotted lines). (B) The same data distributions as in (A), but overlaid with an association model of two *equal* rate limiting steps prior to first passage:

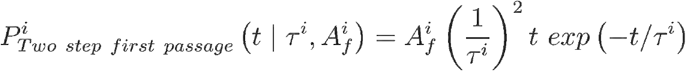

Even ignoring the initial time delay, neither of these models can reproduce the first passage distributions. In general, a lag in the initial association time, like that of Fig. 2B, requires a reaction path with multiple (more than two in this instance) rate limiting steps prior to activation, like that of Eq. 1.

**Supplemental Figure 4.**
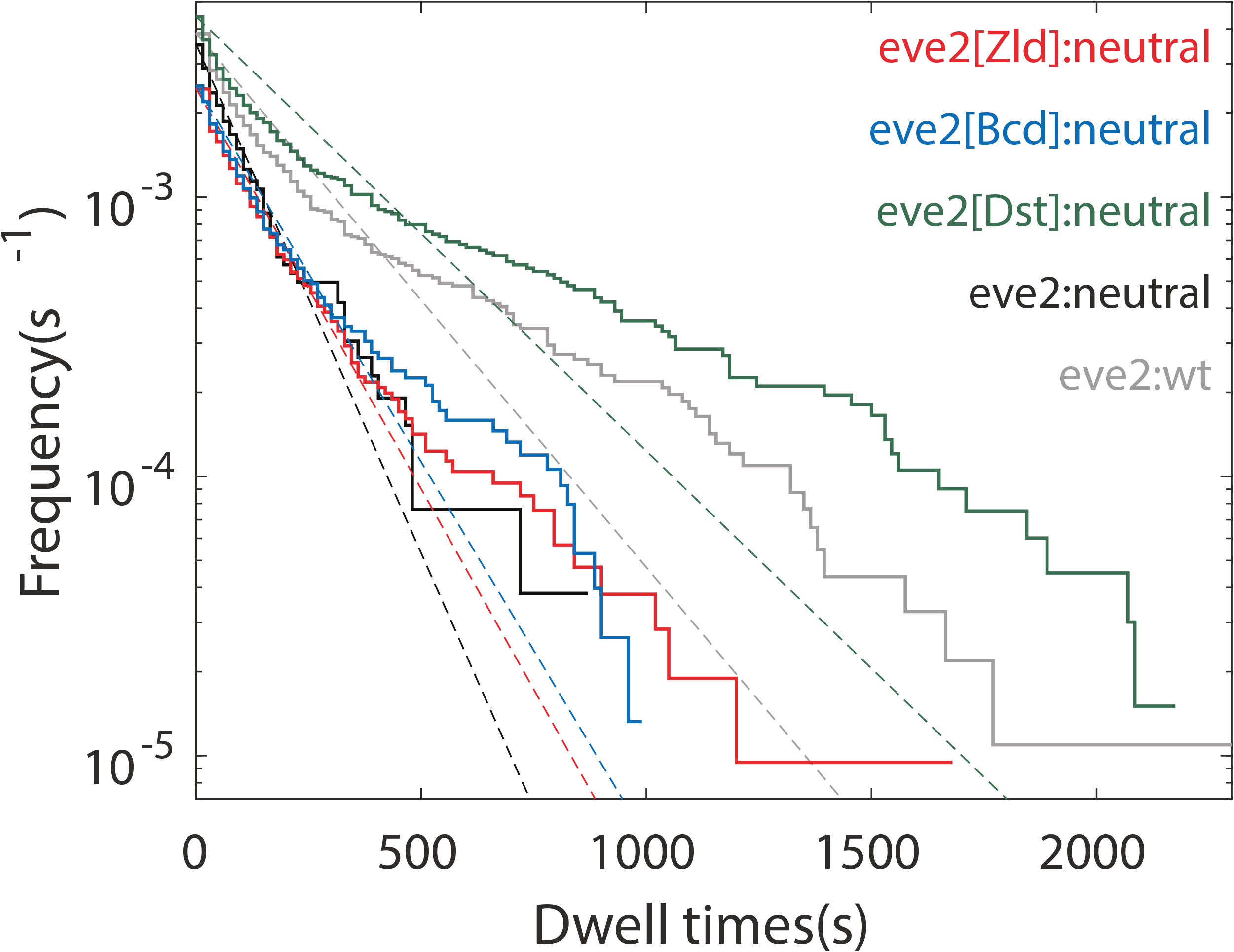
A single characteristic lifetime is insufficient to model active transcription. Cumulative frequency distributions of active transcription lifetimes reproduced from Fig. 3B. For clarity, the shaded error regions have been omitted. In this instance, the data has been overlaid with a single exponential lifetime decay model:

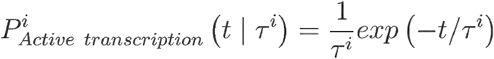

with a single fit parameter, τ^*i*^ (dotted lines). From visual inspection, the distribution predicted by this model is not consistent with the experimental data distribution.

**Supplemental Figure 5.**
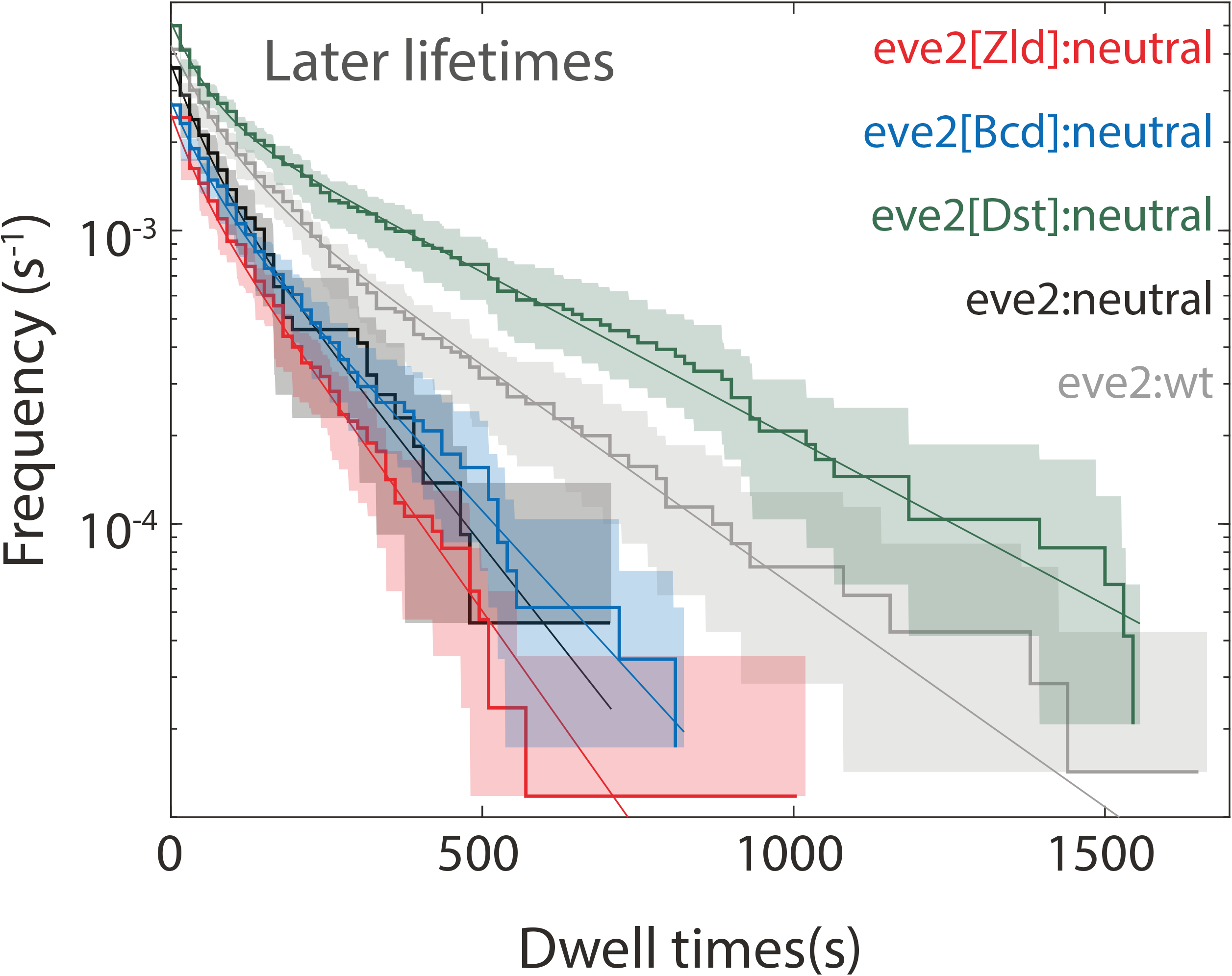
Active transcription kinetics later in NC 14. Cumulative frequency distributions of active transcription lifetimes, as in Fig. 3B, but omitting the 20% of lifetimes that are detected first for each reporter. Put another way, the data shown here combined with the data of Fig. 3C make up the entire distribution shown in Fig. 3B. See Table 2 for parameter values. Shaded regions represent the 90% confidence intervals from bootstrapping methods.

**Supplemental Figure 6.**
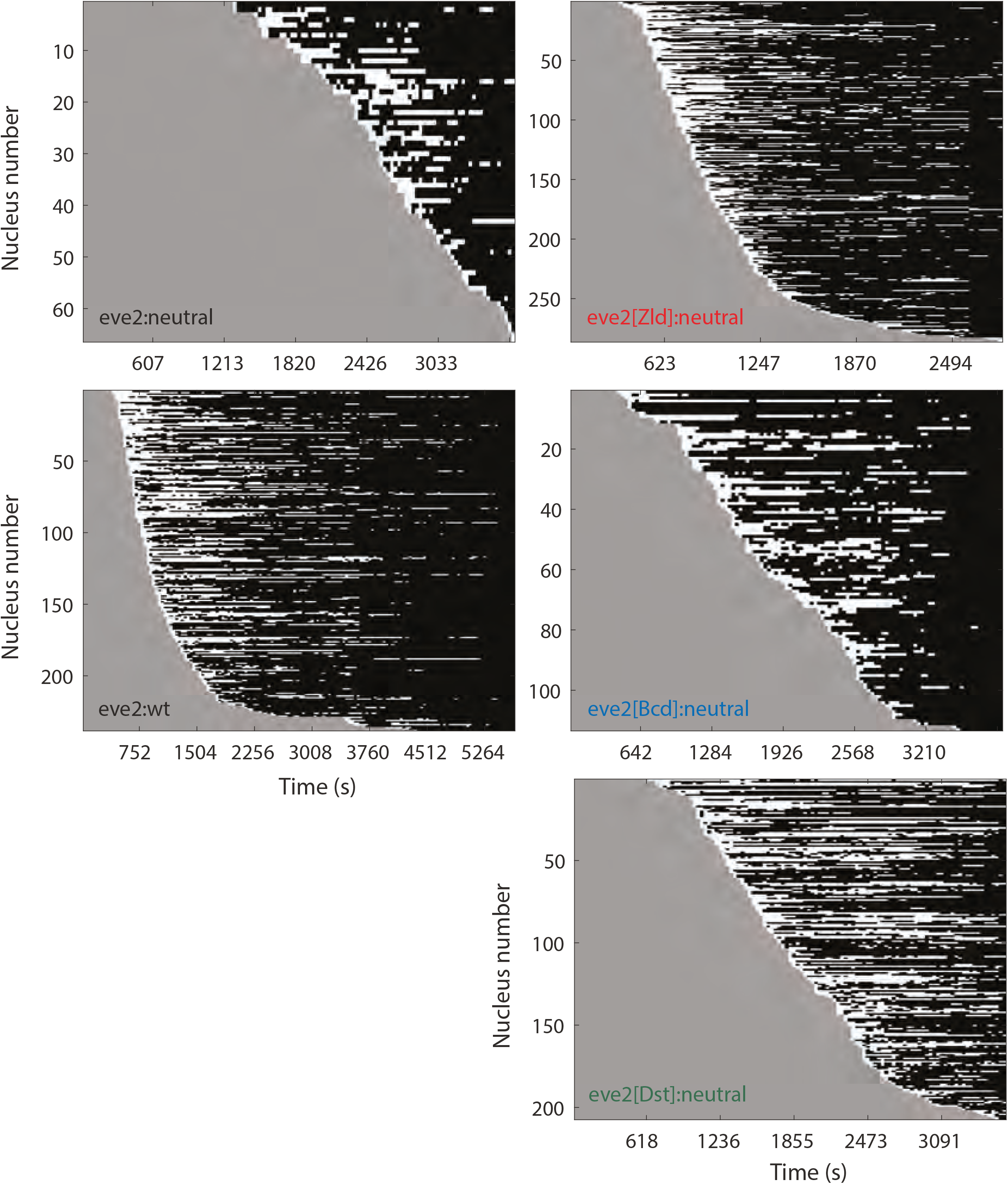
Binarized active transcription time series data for individual nuclei. Raster plots of MCP-GFP transcription signal in every nuclei that displays at least one signal interval. These data are not limited to the center of the stripe. Each row of these plots contains data from a single nucleus over the course of NC 14. Nuclei were sorted by time of first passage into active transcription. Colors indicate first passage interval (gray), active transcription intervals (white), and idle periods (black). The cumulative length of time during which no transcription signal was detected,*L*^*i*^, that was used to compute the mean idle period for each reporter was the sum of the black intervals in each plot (Eq. 3 and Fig. 3).

